# Identifying Causal Subsequent Memory Effects

**DOI:** 10.1101/2021.11.08.467782

**Authors:** David J. Halpern, Shannon Tubridy, Lila Davachi, Todd M. Gureckis

## Abstract

Over 40 years of accumulated research has detailed associations between neuroimaging signals measured during a memory encoding task and later memory performance, across a variety of brain regions, measurement tools, statistical approaches and behavioral tasks. But the interpretation of these Subsequent Memory Effects (SMEs) remains unclear: if the identified signals reflect cognitive and neural mechanisms of memory encoding then the underlying neural activity must be causally related to future memory. However, almost all previous SME analyses do not control for potential confounders of this causal interpretation, such as serial position and item effects. We collect a large fMRI dataset and use a novel experimental design and analysis approach that allows us to statistically adjust for all exogenous confounding variables. We find that, using standard approaches without adjustment, we replicate several univariate and multivariate subsequent memory effects and are able to predict memory performance across people. However, we are unable to identify any signal that reliably predicts subsequent memory after adjusting for confounding variables, bringing into doubt the causal status of these effects. We apply the same approach to subjects’ judgments of learning collected during an encoding period, and show that these behavioral measures of encoding quality do predict memory after adjustments, suggesting that it is possible to measure signals at the time of encoding that reflect causal mechanisms but that existing neuroimaging measures may not have the precision and specificity to do so.

---

What are the neural mechanisms that cause the encoding of lasting memories? The dominant method for identifying successful-encoding-related signals using non-invasive neuroimaging is the subsequent memory paradigm (Paller and Wagner, 2002). This analysis approach compares neuroimaging signals, collected while a subject processes several items, based on later behavioral memory performance (e.g., comparing signals from items that were later remembered to those that were later forgotten). A signal that consistently differs between subsequently remembered and forgotten items suggests a link between the particular underlying activity and memory encoding. These differences, known as subsequent memory effects (SMEs), reliably appear in a number of brain regions, using a variety of memory tasks, imaging technologies and statistical techniques (Kim, 2011; Paller and Wagner, 2002; Xue, 2018). Researchers have theoretically linked particular SMEs to specific latent cognitive or neural mechanisms of memory encoding such as attention, fatigue, representational fidelity, degree of associative binding or match to personal schemas (Sanquist et al., 1980; Wagner, 1998; Davachi et al., 2003; Xue et al., 2010a; Uncapher and Rugg, 2009; Aly and Turk-Browne, 2016; Charest et al., 2014; van Kesteren et al., 2010; Jenkins and Ranganath, 2010; Lohnas et al., 2020; Rugg et al., 2012). In addition, SMEs have for distinguished between multiple cognitive theories of memory encoding and learning (Xue et al., 2010b; Berens et al., 2018) and have been used practically to guide neural stimulation or optimization of learning for improving memory (Ezzyat et al., 2018; Fukuda and Woodman, 2015).

While identifying signals that are associated with memory performance can be of interest itself, claims that the neural activity underlying SMEs reflect encoding mechanisms requires that the activity be *causally* involved in encoding. For activity to be *causal encoding activity*, it means that, if we *were* able to manipulate the activity during the encoding of an event while holding all other external (i.e., non-neural) memory-related factors constant, memory performance would be better, on average, in one condition than the other ^1^. However, identifying causal encoding activity experimentally is exceedingly difficult, if not impossible, because of our limited ability to precisely manipulate neural activity, particularly in humans where there are additional ethical constraints. Therefore our best hope is to learn as much as we can about causal encoding activity from large scale observational data, such as from neuroimaging studies. We argue rather than avoid using causal language altogether to describe the results of observational studies, being upfront about the causal goal can allow us to evaluate to what extent our strategies for identifying causal meet that standard (Weichwald and Peters, 2021; Grosz et al., 2020; Hernán, 2018). In some cases, a simple association, like an SME, can be interpreted causally, such as when there are no confounding variables that affect both the effect (memory performance) and the cause (neural activity). A key difficulty in the causal interpretation of SMEs is that several stimulus-, task- and context-related variables are known to do exactly that. For instance, the concreteness of a word is known to affect both a word’s probability of recognition and recall on a list (Stoke, 1929; Gorman, 1961; Paivio, 1963) as well as neural activity in several brain regions during visual presentation (Binder et al., 2009; Wang et al., 2010). It is possible that some of this neural activity is only involved in word processing and does not directly affect the memory quality. But, in typical sets of word stimuli, signals reflecting this activity will vary with memory performance, and thus be classified as a subsequent memory effect, regardless of their involvement in the encoding process.

Beyond concreteness, stimuli (including images, words and videos) have also been shown to vary in their intrinsic *memorability* (i.e. the average performance for an item in a particular memory task in the population, (Bainbridge, 2019; Baldwin and Runkle, 1967; Isola et al., 2011; Han et al., 2015; Rubin, 1985; Xie et al., 2020)). The concern that confounding variables exist in subsequent memory analyses is not just theoretical: memorability has been shown to drive a significant amount of neural activity in memory tasks during both encoding (Bainbridge et al., 2017) and retrieval (Bainbridge and Rissman, 2018), including in regions typically also associated with SMEs. Therefore, analyses that rely on simply comparing signals between remembered and forgotten trials do not allow for clearly distinguishing between activity that is likely to be causal and activity that merely correlates with memory-predictive features of the stimulus or task.

Does this mean that progress towards uncovering the causal mechanisms for memory encoding from neuroimaging data is hopeless? We believe not. In the social sciences, causal claims are routinely made from observational data, using various approaches to measure and adjust for confounding variables (Angrist and Pischke, 2009; Cochran and Rubin, 1973; Hausman and Taylor, 1981; Pearl, 1995; Winship and Morgan, 1999).

Two recent efforts, Bainbridge et al. (2017), using fMRI with a visual recognition task, and Weidemann and Kahana (2020), using scalp EEG with a verbal free recall task, have attempted to investigate subsequent memory effects while statistically adjusting for item-level memorability and, in the case of Weidemann and Kahana (2020), effects of serial position, a task-level variable that is well known to affect probabilities of recall (Murdock, 1962; Tulving and Arbuckle, 1963). Both papers found that the adjusted subsequent memory effects appeared in a more limited set of brain regions (Bainbridge et al., 2017) and had diminished predictive power (Weidemann and Kahana, 2020). However, as mentioned above, there are many other known effects besides item memorability and serial position that affect the probability of successful encoding such as the distinctiveness or semantic similarity of the item relative to items studied nearby (Aka et al., 2020; Bylinskii et al., 2015; Konkle et al., 2010; von Restorff, 1933). Indeed, the probability of remembering an item in a particular serial position may depend on the item itself. Therefore, even some of the adjusted subsequent memory effects may in fact be confounded by stimulus, task and context effects that drive both causal encoding activity and activity that is unrelated to encoding processes.

A major challenge, then, in identifying observational neuroimaging signals that plausibly reflect causal encoding activity is appropriately measuring all of the confounding variables. In every subsequent memory study we are aware of, presentation is randomized to aid in generalizability. However, from the perspective of dealing with possible confounding factors this presents a combinatorial problem. Estimating the total effect of all exogenous factors that affect memory, including stimulus variables, task variables (such as serial position), contextual effects of a stimulus’ relationship to neighboring stimuli and their possible interactions requires either strong assumptions about the functional form of these effects or an immense amount of data collection to estimate it nonparametrically.

In this study, we circumvent this combinatorial challenge by collecting a unique fMRI dataset of subjects performing a paired-associates verbal memory task where all subjects view the exact same items in the exact same order. This relatively unusual design, inspired by recent studies that measure brain responses across people to a single common experience (e.g., watching the same movie Hasson et al., 2008; Chen et al., 2017), allows us to precisely quantify the total effect of all experimenter controlled exogenous variables in our task.

Through a formal causal analysis (Supporting Information), we can see that this adjustment approach rules out activity that is associated with memory (an SME) but unlikely to be causally involved in encoding processes. Unfortunately, we cannot rule out that activity correlated with memory even after adjusting for confounding is not merely downstream from unmeasured activity that is truly causally involved in encoding (Jones and Kording, 2019). However, we argue that obtaining signals that measure activity downstream of causal activity is still valuable as they as they measure memory-predictive variables that may not be available from behavioral data alone. Therefore, this represents an advance over the unadjusted SME as identified signals provide a more solid foundation for advancing cognitive theory as well as applied goals such as building computeraided systems to improve learning. In addition, it allows for using the relatively cheap observational data to more precisely target plausible sites for expensive stimulation studies. Following the psychometric and epidemiology literature **????**Silva et al. (2002), we define a slightly weaker standard of an *indicator of causal encoding activity* or *ICEA*, that is, activity that is either on the causal pathway to encoding (i.e., causal encoding activity) or downstream from such activity.

Leveraging the above approach, the goal of the current study is to investigate whether several fMRI subsequent memory signals proposed in the literature appear to measure ICEA. While early fMRI studies on subsequent memory effects focused on demonstrating differences in univariate activation (**Activity**)^2^ (Kim, 2011; Paller and Wagner, 2002), subsequent work expanded the types of signals studied to include multivariate patterns (Rissman and Wagner, 2012; Xue, 2018). Deriving these signals involves comparing the spatial pattern of voxel activation across time points or across people (commonly known as representational pattern similarity (Kriegeskorte, 2008; Edelman et al., 1998)) and the ability of these measures to predict memory in a variety of tasks has been extensively documented in the past decade (Xue, 2018). In general, we can categorize these signals into three classes. The pattern similarity of repeated presentations of the same item (Item Pattern Similarity or **IPS**) (Xue et al., 2010b, 2013;

Ward et al., 2013; Visser et al., 2013; Bruett et al., 2020; van den Honert et al., 2016), the pattern similarity of presentations of an item to other items studied in the same encoding period (Global Pattern Similarity or **GPS**) (Bainbridge et al., 2017; Cowan et al., 2020; Davis et al., 2014; Ezzyat and Davachi, 2021; LaRocque et al., 2013; Tompary and Davachi, 2017; Xiao et al., 2016), and the pattern similarity of the same presentation of an item to other participants in a study (Inter-Subject Pattern Correlation or **ISPC**) (Chen et al., 2017; Hasson et al., 2008; Koch et al., 2020). To test these four signal types in a variety of brain regions while avoiding losing power due to large multiple comparisons corrections, we use a predictive modeling approach, testing whether many signals jointly predict recall performance in a regularized regression (Yarkoni and Westfall, 2017).

In sum, our approach to identifying fMRI features that measure ICEA combines a highly controlled memory task which allows for precise measurement of confounding variables with predictive modeling approaches that allow us to increase our power to estimate potentially small associations. Aside from presenting items in the exact same order, our experiment has several other unique features. One particularly interesting application for causal encoding activity might be to track learning of novel information in an educational setting. It may be possible to use measurements of ICEA to adjust teaching plans in a way that will facilitate faster learning, as in automated tutoring systems (e.g. Anderson et al., 2010). We therefore use a novel-language learning task in which participants learn associations between English words and their Lithuanian translations (Nelson and Dunlosky, 1994; Grimaldi et al., 2010). To allow for measuring within-item pattern similarity, our experiment (Fig. 1) consists of presenting 45 paired associates 5 times each during an initial encoding session. This increases overall memory accuracy and the robustness of the item-level measurements. As a comparison to fMRI measures of encoding, we additionally ask participants to provide subjective “judgments of learning” (JOLs), a common behavioral measure related to encoding confidence and metacognition (Arbuckle and Cuddy, 1969; Lovelace, 1984; Nelson and Dunlosky, 1991), for each studied word pair. Finally, participants are tested for their cued recall performance three days after the initial encoding, ensuring that we are measuring signals of durable memory encoding.

**Figure 1:**
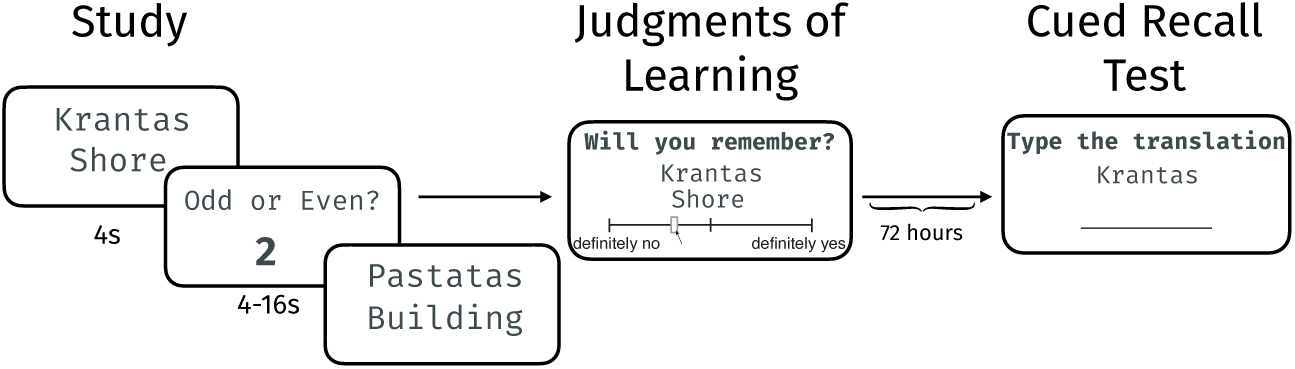
In Phase 1, in the fMRI scanner, subjects first completed a paired associates learning task in which an English word and a Lithuanian word were simultaneously presented on the screen. In between trials, subjects completed an odd-even judgment about a series of numbers in order to prevent rehearsal. After the learning session, subjects made a judgment of learning about each of the paired associates outside of the scanner. In Phase 2, 72 hours after Phase 1, subjects completed a cued recall test in which they were presented with a Lithuanian word and typed in the corresponding English word.

We provide evidence that many previously identified SME signals are able to robustly predict memory formation in our task even in entirely held-out subjects. However, we also show that there is no statistical evidence that these signals reflect neural activity that we can claim is *causally* involved in memory encoding after accounting for possible confounds. Finally, we use the same framework to determine whether participants own judgments of learning are related to ICEA. Repeating our analysis approach, we show that, perhaps surprisingly given the long delay between study and test, it is in fact possible to measure signals (e.g., subjects behavioral responses) at encoding time that do reflect the quality of encoding processes. However, current fMRI indices of memory encoding do not yet have the required level of precision and specificity to measure them reliably.

## Results

### Behavioral Performance

We quantified both the average performance of each of 57 subjects across all 45 word pairs (median = 37%, SD = 15%) and the proportion correct for each word pair across all subjects (median = 33%, SD = 26%). Because of the design of our study, these memorability measures correspond to the proportion correct for a word pair in a particular sequence in the list that every subject in our study observed. The distributions of word pair memorabilities and subject memory abilities are shown in Figure 2. To gain a further sense of how much of the variance in memory performance was explained by the task itself, prior to examining neural activity, we fit a one-parameter item-response theory (IRT) model and examined it’s predictive performance. Specifically, we fit the model *P* (*r*_*ws*_ = 1|*s, w*) = logit^*−*1^(*θ*_*s*_ + *b*_*w*_) with *θ*_*s*_ ∼ *N* (0, *σ*_*θ*_) where *w* indexes Lithuanian-English word pairs, *s* indexes subjects, *r*_*ws*_ is the response (correct or incorrect recall) of each subject to each word pair and *b*_*w*_ and *θ*_*s*_ represent the latent word memorability and subject ability parameters respectively. This model estimates one parameter for each subject which captures their overall ability, and one parameter per word-pair which captures its overall difficulty in the population our subjects were drawn from. We fit this model repeatedly, leaving one subject’s data out and evaluating it’s predictive performance based on the area under the the receiver operating characteristic curve (AUC Green and Swets, 1966; Hanley and McNeil, 1982; Bradley, 1997), computed for the heldout subject. This **IRT** model achieves an average heldout AUC of .72 (Figure 6).

**Figure 2:**
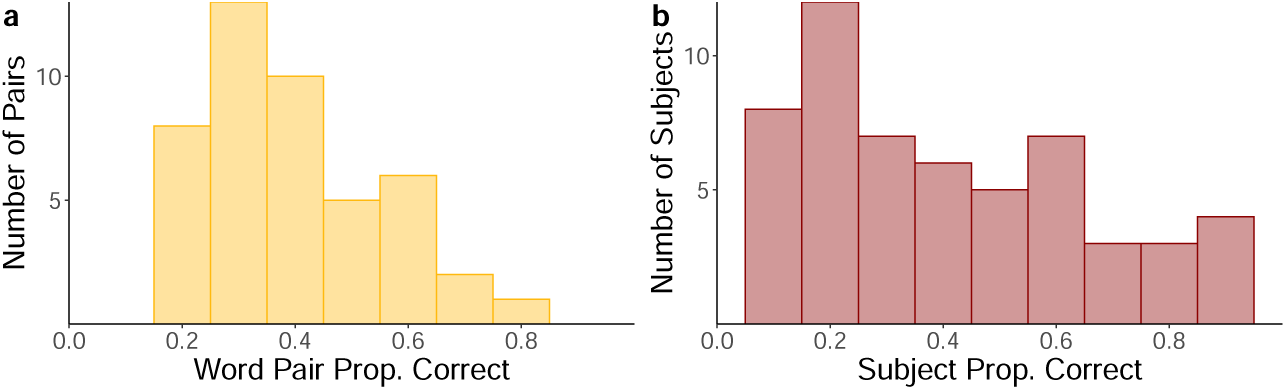
Panel **a** shows the distribution of word pair memorabilities (percent of subjects remembering the pair in a cued recall test) across all 45 word pairs. Panel **b** shows the distribution of subject abilities over all 57 subjects.

### Neuroimaging Analyses

We next attempted to predict memory performance on each individual word pair for each subject from the four types of neuroimaging signals described above, which each define a method for computing predictive features from a set of neuroimaging measurements. Fig. 3 shows how we computed each of the multivariate measures from the neural signals recorded during the task. We use a standard ridge regularized logistic regression approach (Cessie and Houwelingen, 1992), which has commonly been used for studying the neural predictors of memory (e.g. Kuhl et al., 2012; Rissman et al., 2010; Weidemann and Kahana, 2020; Chakravarty et al., 2020), and fit separate models for each of the four features, allowing us to interpret which of the features were associated with successful memory prediction, both without and with adjustments for the memorability of individual word pairs within the context of the list. We focus on linear models here due to their widespread use in the literature and leave evaluating possible non-linear relationships, such as interactions across time and space, to future work.

**Figure 3:**
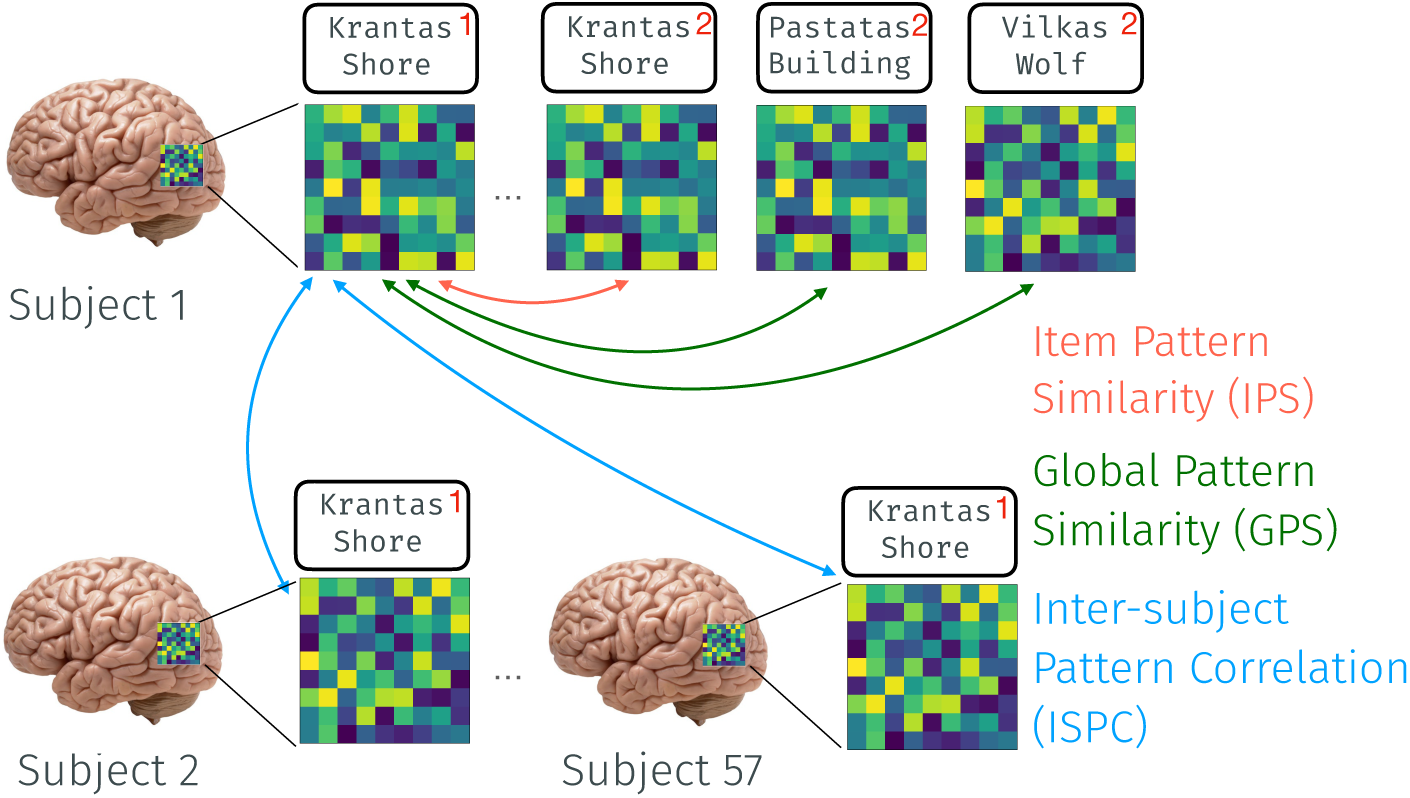
Methods for constructing the three types of multivariate neural features from fMRI signals recorded during encoding. Each word pair in the experiment was presented 5 times and red numbers in the upper right corner indicate the repetition number (but were not present during the actual experiment). Item Pattern Similarity (**IPS**) is computed by comparing two presentations of the same word pair. Global Pattern Similarity (**GPS**) is computed as the mean similarity between a given word pair presentation and presentations of all other words pairs from a different repetition (shown here: repetition 1 and 2). Inter-Subject Pattern Correlation (**ISPC**) is computed by comparing a single presentation of a word pair for one subject to the same presentation of that word pair for all other subjects and averaging the similarities.

To compute the features, we first estimated individual voxel BOLD activation associated with the onset of each study trial with a general linear model (GLM) using the *least squares-single* approach described in (Mumford et al., 2012). Given these maps of activation, we could then compute the four features associated with each trial presentation. Ideally, because regularization down-weights features that are less useful for prediction, we would include every feature in a single model and infer the optimal predictive model. However, in practice, it is well known that including more irrelevant features requires more regularization, limiting predictive performance (Theobald, 1974). In addition, it has long been acknowledged that smoothing can improve the performance of MVPA analyses (Kriegeskorte et al., 2006). This is especially true in the across-subject prediction setting where individual voxels may not be perfectly aligned (Wang et al., 2020; Haxby et al., 2020). Finally, taking averages (or weighting a number of features equally) can have good statistical properties over learning weights for individual features, especially in small datasets (Dawes, 1979; Schmidt, 1971). We therefore describe several approaches below for models that vary in their number of features as well as amount of aggregation prior to computing features, which allows us to test the sensitivity of our approach to these two concerns and strike a balance between model flexibility and sensitivity to small effects.

Because each word pair was presented five times, we considered several approaches to combining the five measurements to predict a single recall test. One approach is to simply include **all** features, one for each study trial, and allow the statistical model to infer their relative importance from the training data, as it may be that measurements taken early on or closer to the end of the study session might be more relevant than others. Another approach is to take the **mean** of each feature across the five presentations. This approach has been used in previous studies with multiple presentations of the same item (e.g. Xue et al., 2010b).

Multivariate neuroimaging signals necessarily involve computing the relationship between several measurements, typically neighboring ones (e.g., grouped via an anatomical map into pre-defined brain regions of interest or using a searchlight analysis (Kriegeskorte et al., 2006)). In order to make our models using univariate activity more comparable in terms of number of features, as well as make them more robust to across-subject deviations, we aggregate voxel activation at the region of interest (ROI) level as well. In our most flexible version of the model, we computed each multivariate feature in each of 100 cortical ROIs from a parcellation of the brain defined by **Schaefer** and colleagues (Schaefer et al., 2018) (Figure 4A). In order to restrict the number of irrelevant features, we test models using only a subset of **targeted** ROIs, selected to reflect the domain knowledge about regions likely to be involved in memory encoding in a verbal learning task (See *Methods* and Figure 4B for the exact definitions). Due to the central role the **hippocampus** is thought to play in memory formation (Scoville and Milner, 1957; Marr, 1971; Norman and O’Reilly, 2003) and because of the difficulty in recording precise fMRI BOLD signal from the hippocampus when using standard imaging sequences for targeting the whole brain (Litman and Davachi, 2008), we test several models including only features computed in the hippocampus. The separate set means that the smaller signal-to-noise ratio in hippocampus BOLD will not prevent its inclusion in the penalized regression models we use for prediction. We test models that include the average activity, pattern similarity and global pattern similarity in each individual’s **whole** hippocampus and also hippocampal **subparts** (left and right, posterior, medial and anterior). In addition, we also select the voxels that are included in all participant’s anatomically-defined hippocampus which allows us to test a classifier based on individual hippocampal **voxel** activity. Finally, we can define a **whole** hippocampus inter-subject pattern correlation feature based on these overlapping voxels (Figure 4C).

**Figure 4:**
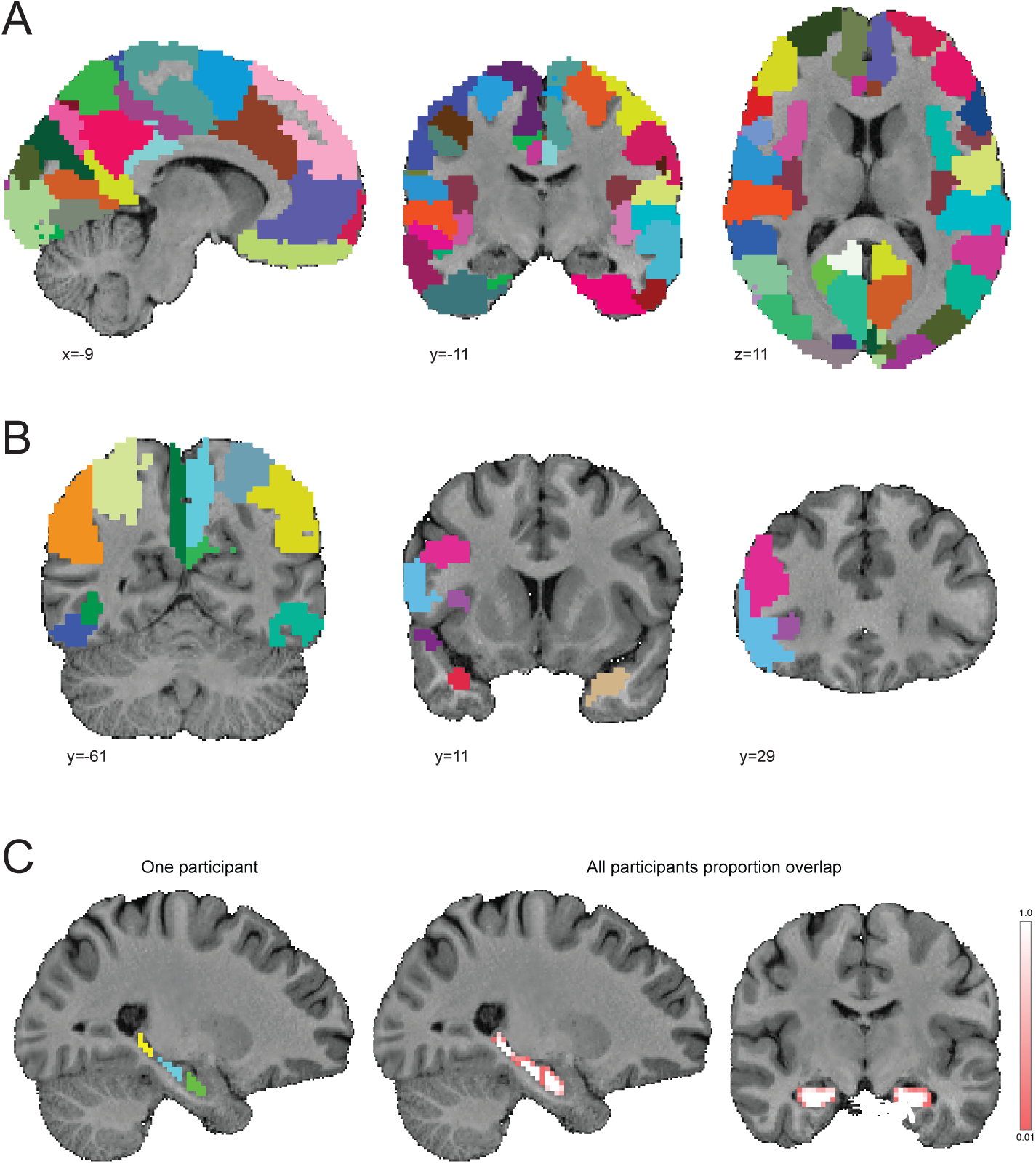
Example location (MNI) and extent of ROIs used for predictions. A) Schaefer et al. (2018) 100 ROI parcellation; B) Example of the targeted ROIs; C) Anatomically defined anterior, mid, and posterior hippocampal ROIs for one participant (left) as well as the proportion of participants having a particular voxel assigned to one of the three hippocampal ROIs (middle and right)

As a final strategy for decreasing sensitivity to errors in voxel alignment, we follow a **whole-brain** strategy, inspired by Rissman et al. (2010), where we treat the average activity in an ROI (using the same set of Schaefer et al. (2018) cortical ROIs as above) as a “voxel” for the purpose of computing similarity across time points in multivariate features.

### Subsequent Memory Models

We first test the ability of these features to predict in a standard subsequent memory setting, that is, predicting memory performance only from features of the neuroimaging signals. These models have the form 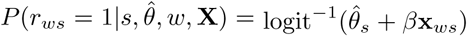 where *w* indexes a word pair, *s* indexes a subject and **X** is one of the sets of neuroimaging signals defined above, normalized within subject. Because this model was fit across subjects, we also included subject-level intercepts to account for overall subject differences in recall performance. To accomplish this, we used the estimates of subject ability from the above-described IRT model 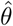. We estimate the predictive power of these models with leave-one-subject-out cross-validation (Stone, 1974), using the area under the ROC curve as a performance metric. Within each training set, we fit these models using a ridge penalty on the *β* parameters and choose the amount of regularization using ten-fold cross-validation. To evaluate the models’ generalization performance, we conduct statistical tests on the held-out AUCs. For the standard models, we compute a one-sample t-test, comparing the model’s performance to the AUC of a random guessing model (.5). In leave-one(-subject)-out cross-validation, the distribution of AUC scores (or any other metric) across holdout sets will in general be correlated because the training set for the classifiers will be largely the same. The independence assumptions of the t-test are therefore violated (Dietterich, 1998). To remedy this, we use a permutation test to estimate the empirical null distribution of paired *t*-statistics when there is no relationship between the fMRI features and recall (Pereira and Botvinick, 2011; Pitman, 1937; Stelzer et al., 2013; Golland and Fischl, 2003).

In Figure 5, we plot the mean and standard error of the AUC estimates in held-out subjects. We show results using both the mean feature across all repetitions as well as using all study block and study-block pairs. Several of our feature/ROI combinations, when aggregated at the mean level, predict memory significantly above chance (.5) based on a permutation test. These include ISPC, GPS and activity models using all Schaefer ROIs as well as when using only the set of Targeted ROIs (M = .56-.63, permuted *p*s *<* .05). Results are overall similar when including each time point separately except that the ISPC models and the Targeted GPS model are no longer significant. Because we are testing so many models it may make sense to control the overall false discovery rate across all models tested (Benjamini and Hochberg, 1995). When we calculate the adjusted *q* values, the Schaefer GPS model including all time points and the Targeted mean GPS model are no longer significant.

**Figure 5:**
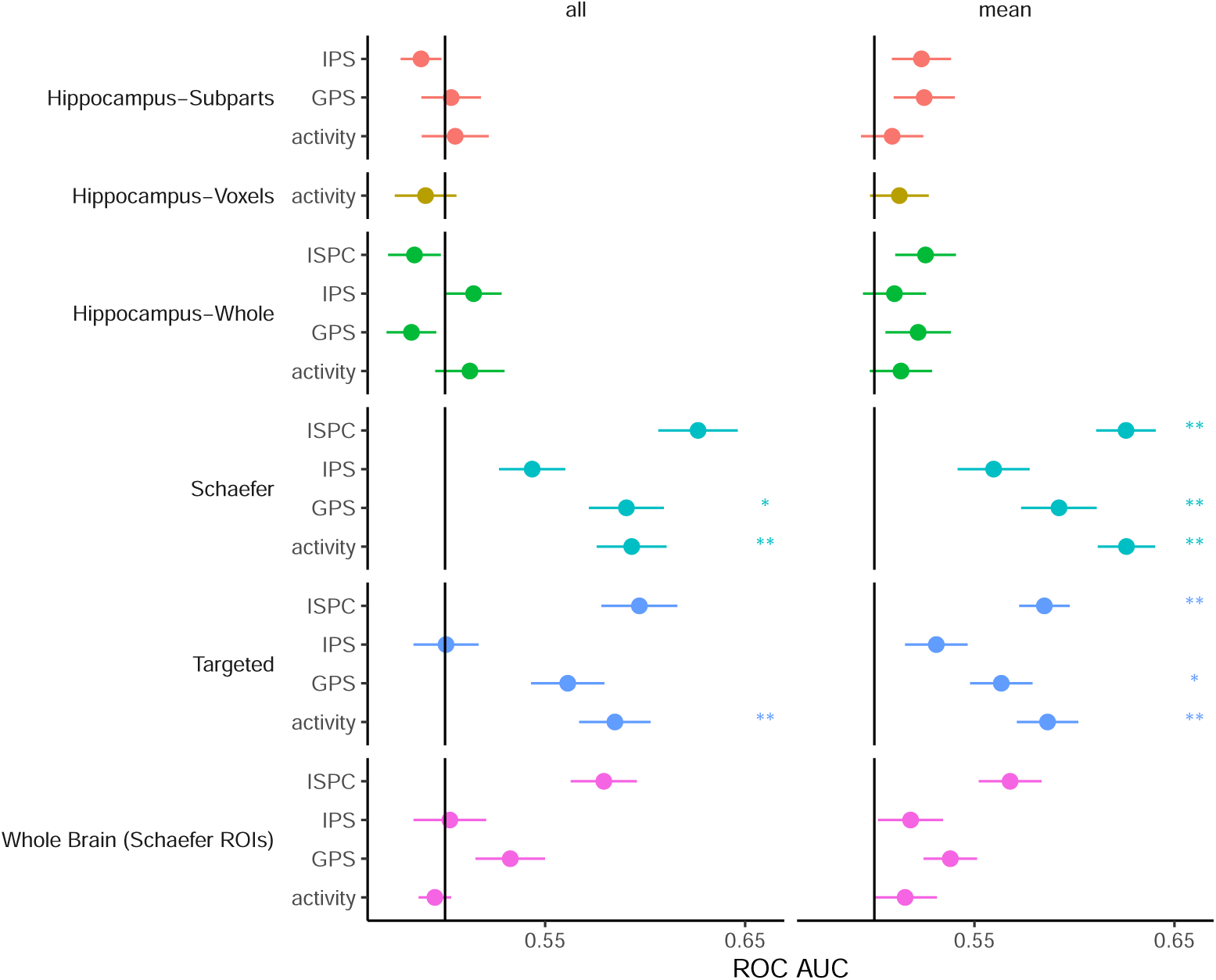
Classifier performance based on the *Standard Subsequent Memory Model*. Each combination of ROIs, features and time-point treatment is plotted separately with definitions of terms found in *Methods. IPS* = Item Pattern Similarity, *GPS* = Global Pattern Similarity, *ISPC* = Intersubject Pattern Correlation. Black line indicates chance AUC (.5) and statistical tests are compared with this baseline. * = *p <* .05 based on a permutation t-test, ** = *q <* .05 after false discovery rate (FDR, Benjamini and Hochberg, 1995) . Error bars reflect the unadjusted standard error of the mean.

**Figure 6:**
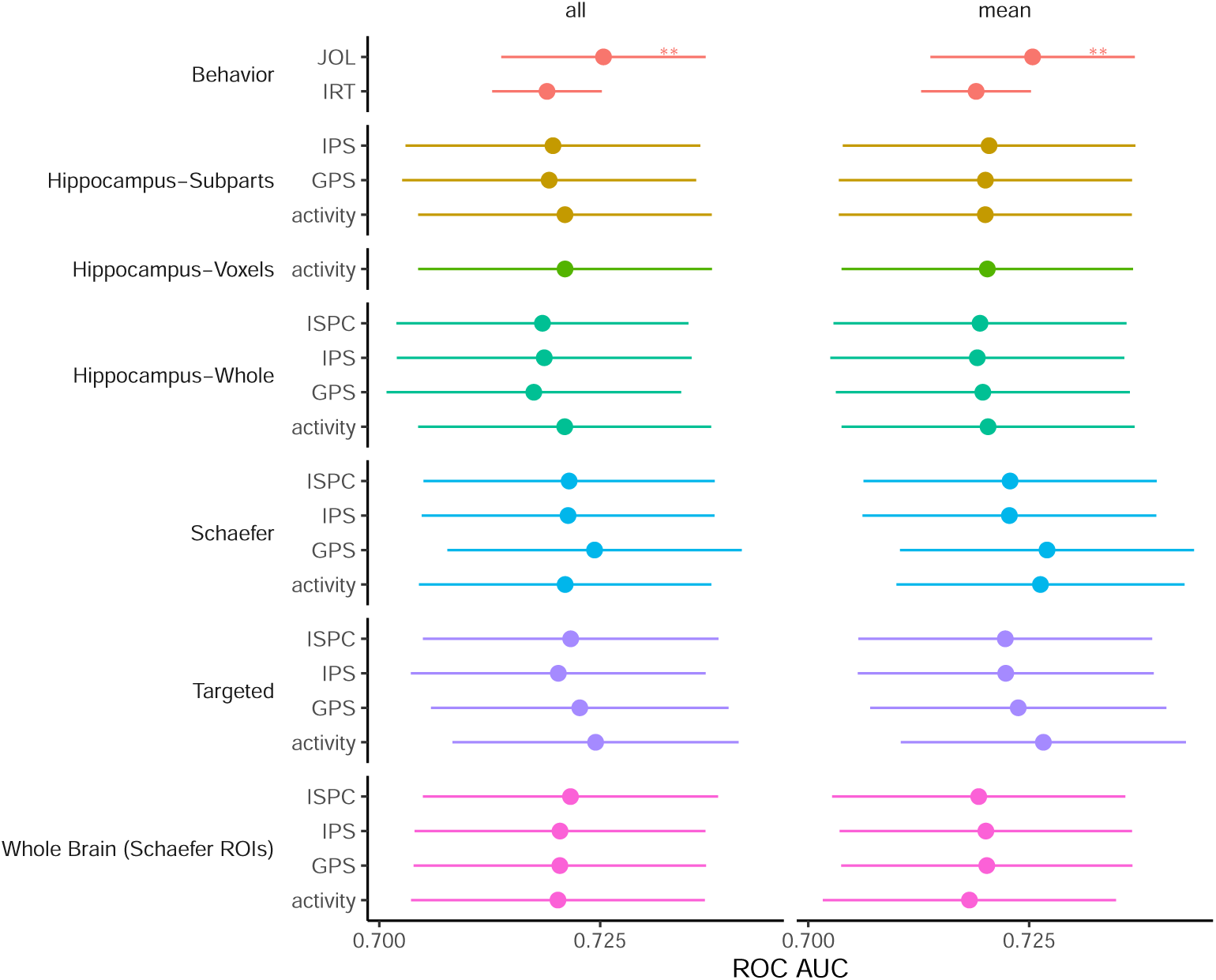
Classifier performance based on the *ICEA Subsequent Memory Model*. Each combination of ROIs, features and time-point treatment is plotted separately with definitions of terms found in *Methods. IPS* = Item Pattern Similarity, *GPS* = Global Pattern Similarity, *ISPC* = Intersubject Pattern Correlation, *JOL* = Judgment of Learning, *IRT* = Item Response Theory model. JOL and IRT are plotted in each column for comparison although they do not differ across columns. * = *p <* .05 based on a permutation t-test, ** = *q <* .05 after an FDR (Benjamini and Hochberg, 1995) adjustment. Error bars reflect the unadjusted standard error of the mean.

Overall, this suggests that several features we tested are able to successfully predict memory across subjects and would normally be classified as subsequent memory effects. In particular, when averaging over all study blocks and including all cortical ROIs, ISPC, GPS and univariate activity could all successfully predict memory in a heldout subject. To our knowledge, this is the first demonstration of across-subject prediction of recall from neuroimaging measures of encoding. This demonstrates that these features of the neuroimaging signal contain a significant amount of information about memory performance. Next, we will test whether there is evidence that the signals reflect activity that is causally involved in encoding or downstream from such activity.

### ICEA Subsequent Memory Models

The IRT model described above estimates the expected performance for each item in the list without considering a subject’s neural signals, essentially estimating the total effect of potential confounding variables (mean AUC = .72, SD =). Given the performance of that model, we can see that the potential confounding factors in a standard memory task already account for a significant amount of the variance in memory behavior.

To estimate ICEA-related subsequent memory effects, we then combine this IRT model with a model for predicting memory from the neuroimaging features. If this model has a stronger relationship with memory performance than the IRT model alone, we argue that this provides evidence that the neuroimaging features reflect ICEA. To do so, we fit a ridge regularized logistic regression that includes a fixed intercept (or offset), which is the linear predictor for each subject and word pair that was estimated previously in the IRT model, i.e.,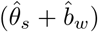. The full model we estimate is therefore: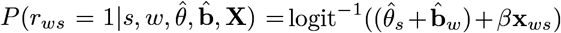, where **x**_*ws*_ is a vector of neural features (e.g., average activity in several ROIs) for subject *s* on word pair *w*. The distribution of heldout AUCs for each model is plotted in figure 6. We compare each model’s heldout AUC performance to the IRT model using a paired *t*-test and using permutations to construct the null distribution. None of our fMRI feature-based classifiers using the mean of the study blocks or all study blocks/study-block pairs perform significantly better than the IRT, even prior to adjustments for multiple comparisons (all *p >* .07). Thus there is no statistical evidence that any of the fMRI features we included in our models are reliably measuring ICEA.

### Judgments of Learning

We can use the same model we employed to test for whether neuroimaging signals measure ICEA with our behavioral measures of judgments of learning. This essentially tests whether, when subjects are asked to give a judgment of learning, they are reading out neural activity that is causally involved in memory or downstream from such activity. The model including judgments of learning (**JOL**) did predict significantly better than than the IRT model, improving the AUC on average by .0128 (*t*_(56)_ = 6.84, *p <* .002).^3^

## Discussion

We found that several univariate and multivariate features of fMRI data proposed in the subsequent memory literature (Kim, 2011; Xue, 2018) could predict cued recall over a long delay in a verbal paired-associates task. To our knowledge, this is the first demonstration of subsequent memory effects in such a task. We also provide, as far as we are aware, the first demonstration that Intersubject Pattern Correlation (ISPC) is a predictor of memory in a task not using video stimuli and that Global Pattern Similarity (GPS) outside of hippocampus predicts memory in a recall task.

However, like in many memory experiments, a significant amount of the variation in the probability of a subject remembering a particular item was explained by the average performance in the population for the specific item in its particular experimental context. A common perspective on subsequent memory effects is that they index “the depth of encoding of the to-be-remembered stimuli” and “determine the efficacy of memory encoding” (Fukuda and Woodman, 2015), suggesting that the underlying activity is causally involved in memory encoding. But because the effects that lead an item to be more likely to be remembered on average might also drive neural activity, the existence of variation in memorability across items confounds a causal interpretation of subsequent memory effects based simply on raw associations. We therefore tested to what extent memory-related fMRI signals were related to variation in recall after adjusting for possible confounding variables such as item memorability (Bainbridge, 2019), serial position (Murdock, 1962) and list composition (Aka et al., 2020). Our analyses, based on comparing predictive models using only behavior with models using behavior and fMRI data combined, showed that the subsequent memory effects we observed were highly correlated with population memorability and did not consistently improve predictions across subjects once the predictive model included the average memorability of an item in a particular context in the population, estimated via a psychometric model. This suggests that the causal interpretation of the activity measured by subsequent memory effects may be less warranted than typically assumed. At least in our data, these analyses suggest that objective characteristics of target items may drive a significant portion of the subsequent memory effect.

Of course, it is entirely possible that our inability to derive a statistically reliable signal of plausible causal encoding activity is simply due to higher variability in our behavioral and neuroimaging data than the typical subsequent memory study. In our experimental design, we did not explicitly manipulate task demands, nor impose a strategy on subjects. In addition, the nature of our task (long delays, associative memory, novel stimuli) may have prevented some processes, perhaps those causally involved in memory formation, from being involved that may typically be measured by subsequent memory effects. However, we argue that the strength of the results using the standard subsequent memory approach, without accounting for confounders, and the results of the ICEA subsequent memory approach when including the judgments of learning as a signal of encoding quality place limits on these alternative explanations. In the following, we discuss several interpretations of these analyses and their implications for future studies of subsequent memory.

One explanation is that much of the variability in memory that could be predicted at encoding is explained by the psychometric model already. For the fMRI features to predict when controlling for the confounding variables, they would have to be reliably indexing causal encoding processes that contradict the prediction from a model based on population averages (e.g., a failure to pay attention to a particular item). In practice, this might be quite rare, especially if participants are focused on the task. In addition, due to the long delay, there is the potential for a lot of variability in memory accessibility between encoding and retrieval due to events that happen in between, limiting the predictability of memory in our task. However, the performance of the model including Judgments of Learning shows that there is a signal (that is in fact consciously available to subjects) that consistently predicts subsequent memory even across items with the same population memorability. This model gives a lower bound on what is explainable at the time of encoding.

A second possible interpretation is that there is too much noise in the fMRI data to hope to find signals comparable to a judgment of learning in a data set our size. However, at 57 subjects, our data set is approximately 2-3 times the size of most data sets in the subsequent memory literature, indicating that future work trying to isolate ICEA may require large data sets and tightly controlled designs.

A third possible interpretation is that there are indeed signals of ICEA that exist in fMRI data but they are highly variable across people, perhaps due to a lack of precise voxel alignment or differently shaped brains causing variation in the signal-to-noise characteristics. If this were the case, it may be that our attempt to do across-subject prediction was doomed from the start. However, we point to the success of our associative subsequent memory models (without adjusting for confounding variables) as evidence that these issues do not prevent any across-subject prediction in our data. In addition, several of our features made efforts to circumvent this difficulty by a) aggregating data at higher levels (e.g. the whole brain analyses treating each ROI as a “voxel”), b) using hippocampal ROI definitions based on individual subjects’ anatomy and c) using multivariate measures like pattern similarity that are less sensitive to perfect alignment across subjects. However, none of these features seem to have succeeded in improving classifier accuracy significantly beyond the IRT model.

A fourth possible interpretation is that the actual cognitive strategies used for learning (especially in an undirected, intentional encoding task like ours) are highly variable across people. For instance, it may be that high-memorability word pairs are easier to sound out, resulting in greater activity in phonological regions on average. But if sounding out is only an effective strategy for some people, activity in a particular phonological region may not consistently predict memory. Across-subject prediction differs from the usual two-level within-subject paradigm that is typical of MVPA studies in that it is more sensitive to small magnitude, low variability effects but less sensitive to high magnitude, high variability effects (Wang et al., 2020). If the measurement of encoding processes is more like the latter scenario, this suggests (like above) that future studies of subsequent memory will require larger data sets that allow for understanding groups of subjects who use similar encoding strategies.

Overall, it is certainly possible that our study was limited by a low neuroimaging signal-to-noise ratio and high across-subject variability in both the fMRI BOLD measurement and the cognitive processes involved in encoding themselves. However, we argue that the success of the JOLs as a predictor of memory after adjusting for confounding and the existence of successful memory prediction from fMRI in the absence of item and task variables suggest that this can’t be the complete story.

## Conclusion

In the forty years since the first subsequent memory effect was reported (Sanquist et al., 1980), much has been learned about the neural signals associated with memory. A vast literature maps the various uni- and multi-variate signals across the brain, along with proposed cognitive computations being implemented by the underlying neural activity. In addition, many signals have supporting evidence from animal and lesion studies. This paper contributes to this literature by documenting, for the first time, several aspects of subsequent memory effects in a novel language-learning task. We show that many features proposed to be relevant for other types of memory tasks such as Intersubject Pattern Correlation and Global Pattern Similarity also apply here. However, the exact interpretation of these subsequent memory effects remains ambiguous, and the original hope of identifying the causal mechanisms of memory encoding has not been fully realized. Here we defined a framework for making progress towards identifying (Indicators of) Causal Encoding Activity from neuroimaging signals. Based on this formulation, we attempted to precisely control for all exogenous effects including stimulus, presentation-order and overall list context effects that have been documented to influence the probability of recall in order to see whether we can extract signals from fMRI that are still associated with subsequent memory performance, reflecting activity that is either causally involved in memory encoding or downstream form such activity. While we were not able to identify such signals, we suggest that future work leverage the techniques of this paper, novel classifier analyses and tight control of memorability, with larger datasets, different tasks and new features of the fMRI signal. In several ways, our own design could be improved, for instance, by using items with more similar memorability scores (so that more variance can be explained by the neural signals), by including a movie-watching portion of the scanner task that would allow for the use of better alignment tools (Haxby et al., 2011, 2020), or by using a task such as recognition memory that would allow for more trials to be collected per participant. Despite these qualifications, we think that these results suggest that interpretations of classic subsequent memory results may not be as straightforward as commonly assumed and lay out key questions for cognitive neuroscientists of memory to address in the future.

## Materials and Methods

### Participants

Sixty-nine participants were recruited from using electronic advertisements hosted by New York University and accessible to the broader community. We obtained written informed consent prior to conducting the study, in compliance with the New York University Institutional Review Board. All participants self reported to be between 18-35 years old, had normal or corrected-to-normal vision, spoke English fluently and did not speak Lithuanian or a related language.

Eleven participants’ data were excluded from analysis based on MRI data issues: two participants’ MRI sessions were conducted with incorrect scanning parameters; six participants were excluded based on excessive motion and other image quality issues; three participants requested early termination of the experiment due to discomfort in the scanner (and did not complete the recall test). We also excluded one of the participants who recorded 100% correct in the recall test since their held-out AUC would be undefined, resulting in a final data set of fifty-seven participants

### Behavioral Task

Based on a normed set of Lithuanian-English words, we selected 45 translation pairs with a range of difficulties (Grimaldi et al., 2010) for use in our cued recall task, also described in Figure 1. All participants first completed a study phase inside the fMRI scanner where they saw the translation pairs presented one at a time for 4 seconds each with a variable duration inter-trial interval (randomly chosen between 4 and 16 seconds). During this inter-trial interval, in order to prevent rehearsal, participants made judgments about whether each of a series of numbers were odd or even.

Words were presented on a computer screen with the Lithuanian word at the top of the screen and the English translation underneath. Each word pair was presented five times and no pair was presented for the *n*th repetition until all words had *n* − 1 presentations. Importantly, and in contrast to many psychology studies on the subsequent memory effect, all participants see the same sequence of study items. At the expense of generalizability, this allowed us to precisely quantify the average recall performance for a word pair in a specific context and accounting for any interactions between items. The order of the word pairs was selected as follows: the 45 words were grouped into groups of five words each. On each repetition the order of the five words were shuffled but all five words appeared before the next group of five were presented. Immediately following the study session participants completed a second section outside the scanner that involved making judgments of learning (JOLs (Nelson and Dunlosky, 1991)): for each pair participants were presented with the Lithuanian and English word and used the computer mouse to indicate on a scale of 0-100 how likely they were to remember the association in three days. Participants had up to twelve seconds to respond and the response was coded as missing if this deadline wasn’t met (in practice, all participants gave all JOL ratings within the allotted time).

Participants were then asked to complete a recall test approximately 72 after the initial scan session. The first 44 of the subjects completed this in the lab and the final 13 subjects completed the second session at home, via an online version of the task to ease the task of scheduling sessions with a 72 hour delay between. Participants saw a Lithuanian word presented on the screen and had to type the associated English word. A trial was coded as correct if participants typed the correct English word (allowing for typographic errors) and all other responses were incorrect. Like the judgments of learning, this task was completed outside the scanner.

### fMRI Data Acquisition

MRI data were acquired on a 3 Tesla Siemens Prisma scanner. Anatomical images were collected using a T1-weighted MPRAGE high-resolution sequence (0.8mm isotropic voxels). Five runs of whole brain BOLD functional data were acquired using an EPI sequence (2.5mm isotropic voxels; TR=1s; TE=35ms; Multiband factor=4; Phase Encoding: Anterior-Posterior). An additional pair of non-accelerated EPI scans with opposing forward (anterior-posterior) and reverse (posterior-anterior) phase encoding relative to the primary functional data were collected for distortion correction during preprocessing (2.5mm isotropic voxels; TR=4.496; TE=45.6). The slice position and orientation for these scans matched those of the BOLD data collected during the behavioral task.

### fMRI Preprocessing

Results included in this manuscript come from preprocessing performed using fMRIPprep 1.2.6-1 (Esteban et al., 2019), which is based on Nipype 1.1.7 (Gorgolewski et al., 2011).

#### Anatomical data preprocessing

The T1w anatomical scans were corrected for intensity non-uniformity (INU) using `N4BiasFieldCorrection` (ANTs 2.2.0, Tustison et al., 2010). A T1w-reference map was computed (after INU-correction) using `mri_robust_template` (FreeSurfer 6.0.1, Reuter et al., 2010). The T1w-reference was then skull-stripped using `antsBrainExtraction.sh` (ANTs 2.2.0), using OASIS as target template. Brain surfaces were reconstructed using `recon-all` (FreeSurfer 6.0.1, Dale et al., 1999), and the brain mask estimated previously was refined with a custom variation of the method to reconcile ANTs-derived and FreeSurfer-derived segmentations of the cortical gray-matter of Mindboggle (Klein et al., 2017). Spatial normalization to the ICBM 152 Nonlinear Asymmetrical template version 2009c (Fonov et al., 2011) was performed through nonlinear registration with `antsRegistration` (ANTs 2.2.0, Avants et al., 2011)), using brain-extracted versions of both T1w volume and template. Brain tissue segmentation of cerebrospinal fluid (CSF), white-matter (WM) and gray-matter (GM) was performed on the brain-extracted T1w using `fast` (FSL 5.0.9 Zhang et al., 2001).

#### Functional data preprocessing

For each of the five BOLD runs collected per subject (across all tasks and sessions), the following preprocessing was performed. First, a reference volume and its skull-stripped version were generated using a custom methodology of *fMRIPrep*. A deformation field to correct for susceptibility distortions was estimated based on two echo-planar imaging (EPI) references with opposing phase-encoding directions, using `3dQwarp` (AFNI 20160207, Cox, 1997). Based on the estimated susceptibility distortion, an unwarped BOLD reference was calculated for a more accurate co-registration with the anatomical reference.

The BOLD reference was then co-registered to the T1w reference using `bbregister` (FreeSurfer) which implements boundary-based registration (Greve and Fischl, 2009). Co-registration was configured with nine degrees of freedom to account for distortions remaining in the BOLD reference. Head-motion parameters with respect to the BOLD reference (transformation matrices, and six corresponding rotation and translation parameters) are estimated before any spatiotemporal filtering using `mcflirt` (FSL 5.0.9, Jenkinson et al., 2002). The BOLD time-series were resampled onto their original, native space by applying a single, composite transform to correct for head-motion and susceptibility distortions. These resampled BOLD time-series will be referred to as preprocessed BOLD in original space, or just preprocessed BOLD. The BOLD time-series were resampled to MNI152NLin2009cAsym standard space, generating a pre-processed BOLD run in MNI152NLin2009cAsym space.

Several confounding time-series were calculated based on the preprocessed BOLD: framewise displacement (FD), DVARS and three region-wise global signals. FD and DVARS are calculated for each functional run, both using their implementations in Nipype (following the definitions by Power et al., 2014). The three global signals are extracted within the CSF, the WM, and the whole-brain masks. Additionally, a set of physiological regressors were extracted to allow for component-based noise correction (CompCor, Behzadi et al., 2007). Principal components are estimated after high-pass filtering the preprocessed BOLD time-series (using a discrete cosine filter with 128s cut-off) for the two CompCor variants: temporal (tCompCor) and anatomical (aCompCor). Six tCom-pCor components are then calculated from the top 5% variable voxels within a mask covering the subcortical regions. This subcortical mask is obtained by heavily eroding the brain mask, which ensures it does not include cortical GM regions. For aCompCor, six components are calculated within the intersection of the aforementioned mask and the union of CSF and WM masks calculated in T1w space, after their projection to the native space of each functional run (using the inverse BOLD-to-T1w transformation). The head-motion estimates calculated in the correction step were also placed within the corresponding confounds file. All resamplings can be performed with a single interpolation step by composing all the pertinent transformations (i.e. head-motion transform matrices, susceptibility distortion correction, and co-registrations to anatomical and template spaces). Gridded (volumetric) resamplings were performed using `antsApplyTransforms` (ANTs), configured with Lanczos interpolation to minimize the smoothing effects of other kernels (Lanczos, 1964). Non-gridded (surface) resamplings were performed using `mri_vol2surf` (FreeSurfer). Preprocessed BOLD data were smoothed to 6mm FWHM using AFNI `3dBlurToFWHM`^4^.

#### Regions of interest (ROIs)

##### Gray matter and hippocampus anatomical masks

Analysis of the functional data was restricted to voxels encompassed by a binary mask computed from the probabilistic gray matter estimates generated during the Freesurfer anatomical preprocessing combined with anatomically defined voxels in individual hippocampi. For each participant, the probabilistic gray matter mask was binarized with threshold of 0.2 and intersected with bilateral hippocampal voxels as estimated using FSL `FLIRT` implemented in Nipype (Jenkinson and Smith, 2001; Jenkinson et al., 2002). The combined gray matter/hippocampus mask was resliced to functional resolution, smoothed with an 8mm FWHM kernel to be liberal in voxel inclusion (we opted to include some potentially non-gray matter voxels that could be ignored in downstream analyses rather than being overly strict and excluding relevant signal) and then re-binarized with a threshold of 0.2.

##### Whole brain cortical ROIs

For whole brain cortical analysis, voxels were aggregated into ROIs provided by (Schaefer et al., 2018). To strike a balance between regional specificity and total number of cortical ROIs we used the intersection of the gray matter mask defined during preprocessing and the 100 parcel, 7 network atlas from Schaefer et al. (2018)^5^ after reslicing to 2.5*mm*^3^.

##### Targeted ROIs

We defined a targeted set of nineteen ROIs on the basis of prior expectations of these regions’ potential engagement in a learning task involving verbal materials. We selected left and right perirhinal cortex (PRC) and anterior temporal lobe (ATL) ROIs due to putative roles for these regions in processing conceptual information (Martin et al., 2018). In addition, we selected regions highly likely to be involved in memory formation (Kim, 2011) and in processing concrete nouns (a category which includes all of the word pairs in our stimulus set) (Chen et al., 2019). These ROIs included: two separate areas in left anterior insula, left and right lateral occipital cortex, left and right dorsal parietal, left and right lateral parietal, left and right medial parietal (precuneus), left and right retrosplenial cortex, an upper and a lower portion portion of left ventrolateral prefrontal cortex (VLPFC), and visual word form area (VWFA; left hemisphere only). This set of targeted regions listed above were defined from a variety of sources. The PRC regions were defined using the anterior portion (MNI *y >*−21) of the PRC ROIs provided by (Ritchey et al., 2015) as downloaded from Neurovault^6^ which overlapped with the PRC center coordinates reported by Martin et al. (2018). The ATL ROIs were defined as the Schaefer et al. (2018) ROIs encompassing the peak voxels reported by Martin et al. (2018) (Schaefer et al. (2018) 600 parcellation ROIs 191 and 492).

The remaining targeted ROIs were defined by identifying voxel groupings from Schaefer et al. (2018) which encompassed other cortical regions expected to be involved in memory formation and/or processing of verbal materials.

#### Single trial activation estimates

Individual voxel BOLD activation on each study trial was estimated with a general linear model (GLM) as implemented in SPM12 (Friston et al., 1994)^7^ using the *least squares-single* approach described in Mumford et al. (2012). For each study trial, a voxel-wise regression was carried out with one predictor containing a four second boxcar covering the timepoints corresponding to the to-be-estimated trial, another predictor containing four second boxcars covering the timepoints for all of the remaining trials, and fourteen additional predictors containing a variety of other potential confounds and nuisance signals as estimated during preprocessing (Power et al., 2014). Prior to estimation, the vectors of single-trial and all-other trials were convolved with the canonical SPM double gamma hemodynamic response function (HRF) along with its temporal and dispersion derivatives (Lindquist et al., 2009). We then estimated regression coefficients for each voxel using ordinary least squares and the procedure was repeated for each study trial for each participant. The resulting coefficients for the single-trial predictors were taken as summaries for the activation for each voxel on each trial.

#### fMRI feature estimates

After initial fMRI preprocessing, additional steps to create the fMRI features for predictive models and reshape data were conducted using custom code in `python 3.7.7` using the packages `pandas` (McKinney, 2010) and `numpy` (van der Walt et al., 2011) and R 4.0.2 using the package collections `tidyverse` (Wickham et al., 2019) and `tidymodels` (Kuhn and Wickham, 2020).

Multivariate patterns were derived from the single trial estimates for each voxel in a particular ROI. While several distance metrics have been proposed (Bobadilla-Suarez et al., 2019; Walther et al., 2016), in order to remain faithful to the proposed features, we used Pearson’s *r*, which is the metric used by nearly every paper in the subsequent memory literature (e.g. LaRocque et al., 2013; Xue et al., 2010b; Davis et al., 2014). In addition, for inclusion in the model, all of the features are Fisher-*z* transformed (Fisher, 1915) which is often used when aggregating correlations since it makes the sampling distribution approximately normal, which is slightly better for use in the regularized regression models we describe in the following section.

### Predictive Modeling

#### IRT Model

We estimate this model using maximum likelihood, as implemented in the R package `lme4` (Bates et al., 2015). Like the models involving fMRI data, these models were trained in a leave-one-subject-out manner. When predictions are made using these models for a heldout subject, *θ*_*s*_ was set to 0 where *s* indexed the heldout subject.

#### Regularized Regression Models

To estimate the models, we use cyclical coordinate descent as implemented in the `glmnet` package (Friedman et al., 2010) in R. We estimate the predictive power of our models with leave-one-subject out cross-validation (Stone, 1974), using the area under the ROC curve as a performance metric. We choose the size of the penalty on the *β* parameters (*λ*) using nested 10-fold cross-validation. We choose a set of *λ* hyperparameters to try using the default algorithm in `glmnet`: first *λ*_max_ is chosen such that a model fit with that *λ* on all of the data will have all parameter estimates near 0, using a formula from Friedman et al. (2010). Then 100 parameters are chosen on a log scale from *λ*_max_ to 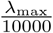. The model is estimated for each *λ* and the *λ* that maximizes the log likelihood of the held-out fold is chosen for evaluation. All of our fMRI-derived features are standardized (by their mean and variance in the training fold) before being included in the model.

### Evaluation

To evaluate the models’ generalization performance, we conduct statistical tests on the held-out AUCs. For the standard models, we compute a one-sample t-test, comparing the model’s performance to the AUC of a random guessing model (.5). AUC is a useful metric here because it is not affected by the base rates of remembering and forgetting. For the main test, we compare the ICEA Subsequent Memory Models to the behavioral model using a paired t-test. To give the strongest possible chance to the MRI features, for all models, we use a *one*-sided test of whether the AUC in the ICEA Subsequent Memory Model is greater. In leave-one(-subject)-out cross-validation, the distribution of AUC scores (or any other metric) across holdout sets will in general be correlated because the training set for the classifiers will be largely the same. The independence assumptions of the t-test are therefore violated (Dietterich, 1998). To remedy this, we use a permutation test to estimate the empirical null distribution of paired *t*-statistics when there is no relationship between the fMRI features and recall (Pereira and Botvinick, 2011; Pitman, 1937; Stelzer et al., 2013; Golland and Fischl, 2003). To do so, we shuffle the pairing between the fMRI features and the Lithuanian-English word pair within subject. This allows us to maintain the correlation between fMRI features, as well as the distribution of word-pair population memorabilities and subject recall abilities. Crucially, this also means that *IRT* does not change its predictions so the residual variance to be explained remains the same across permuted datasets. When shuffling the data, we also maintain the structure of the sequence of items such that the same five items appear in each group of five serial positions of each study block. We created 500 permuted datasets, applied to same classification models and computed a paired *t*-statistic for the permutation test.

## Data Availability

All data needed to reproduce the results in this paper will be made available on a public repository upon publication.

## Code Availability

All code needed to reproduce the results in this paper will be made available on a public repository upon publication.

## Acknowledgements

We thank Hong Yu Wang, Camille Gasser, and Steven Mikal for helpful assistance with stimulus development, scanning, and data processing. We thank Daniel Schonhaut, Noa Herz, John Sakon and Michael Kahana for helpful comments on a previous draft. This work was supported by NSF grant DRL-1631436 and seed funds from the NYU Dean for Science.

## Supporting Information Text

This appendix lays out a causal graphical model approach to understand how causal encoding activity can be identified (Pearl, 1995).

## Causal Models

### Concreteness Model

We first imagine, as in the main text, the situation if the only variable that causally impacted memory (through neural activity) was concreteness. In this model, shown graphically in figure S1, we assume that the concreteness of a stimulus drives neural activity and some of that activity, labeled Causal Encoding Activity, causes the probability of later memory to increase or decrease. This is downstream from concreteness, and can presumably be thought of as something like “perceived concreteness.” Both Causal Encoding Activity and Non-causal Encoding Activity may underlie any particular fMRI signal we are interested in so the key question is how to identify fMRI signals that measure Causal Encoding Activity, or in other words, signals where the red arrow in figure S1 exists. One useful thing we can derive from graphs, using the logic laid out by Pearl (1995), are the implied (conditional) independences. This logic provides easy algorithms for checking dependencies in graphs and they are implemented in packages like `dagitty` in R (Textor et al., 2017). In the graph without the red arrow, when we condition on Concreteness, the fMRI signal will be independent of Memory. This suggests that we can distinguish fMRI signals that measure Causal Encoding Activity from those that do not by testing whether fMRI signals that are not independent from Memory after conditioning on Concreteness.

### Endogenous-Exogenous Model

Following Weidemann and Kahana (2020), we divide up the factors affecting memory into endogenous and exogenous factors. Exogenous factors represent a group of variables that are in principle under experimenter control because they involve features of the item presented or features of the way in which items are presented. Many such features have been extensively documented in the psychological literature as having effects on memory performance. Since these are observable, through the method we describe in the paper, we can simply replace Concreteness in Fig. S1 with the effects of all Exogenous variables as in Fig. S2. In this case, because we can block the path from the non-causal neural activity to memory, we can use our approach to identify causal encoding activity.

In contrast, endogenous features represent ongoing latent cognitive or neural processes that are known to affect memory such as fluctuations in attention or fatigue, or their neural correlates. These processes are necessarily latent and it is difficult to obtain precise measures of them from behavior alone. Figure S3 shows a causal graph in which both sets of variables influence memory. The endogenous variables are represented as a dotted circle because we do not measure them. What becomes clear from this graph is that it is now complicated to separate out the causal activity from the non-causal. This is because if some endogenous neural activity prior to the trial caused both causal and non-causal neural activity to change, without controlling for the endogenous factors, both sources of activity would apper to be correlated with memory.

To clarify exactly which non-causal neural activity will appear correlated with memory, we expand our model further as in Fig. S4. Here, we can separate our endogenous variables into causal and non-causal. Causal endogenous variables represent variables that cause causal encoding activity (and indirectly, memory performance) while non-causal endogenous variables do not. Only signals that reflect causal neural activity or activity downstream from causal neural activity will appear correlated with memory after controlling for the exogenous variables. Therefore, we call these *indicators* of causal encoding activity. As mentioned in the main text, these signals are still valuable to discover as they provide measurements of causal endogenous variables, which may be useful in constructing theories. In addition, because this is a subset of all non-causal encoding activity, this approach restricts the set of signals of interest for future studies.

**Figure S1:**
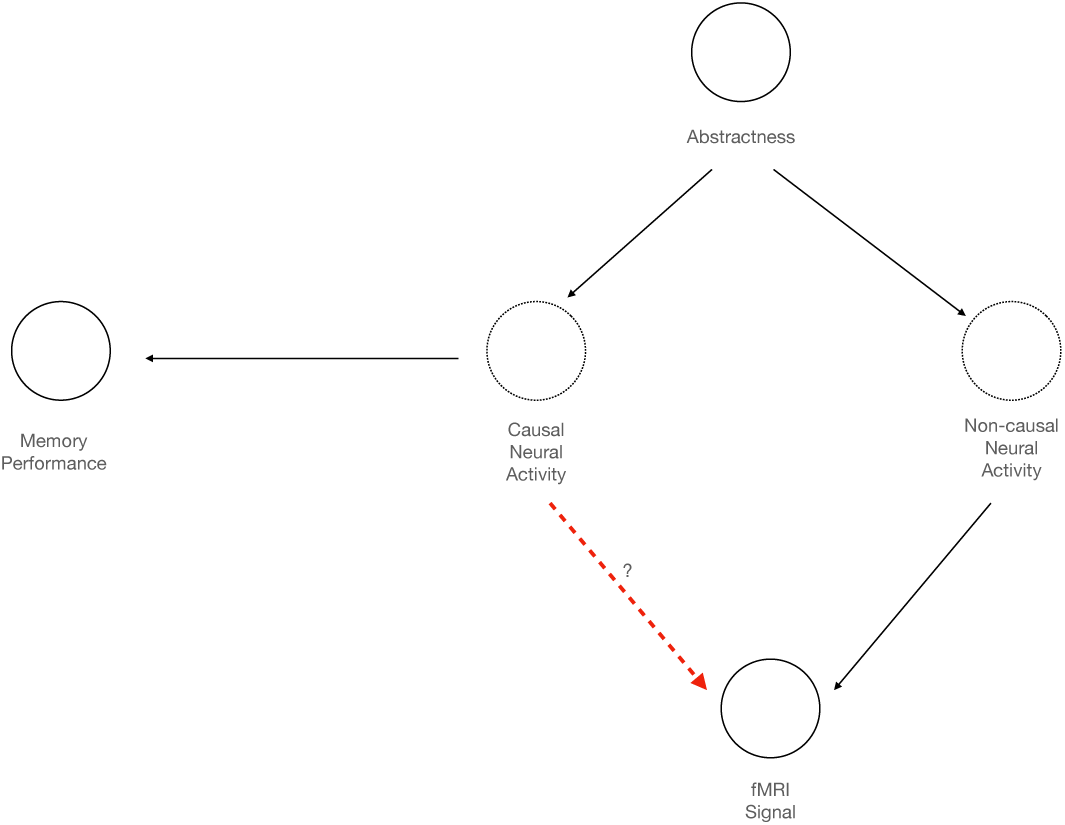
A directed acyclic graph representing a causal model for memory if abstractness was the only variable that influenced memory. Solid circles indicate observed variables while dotted circles indicate unobserved.

**Figure S2:**
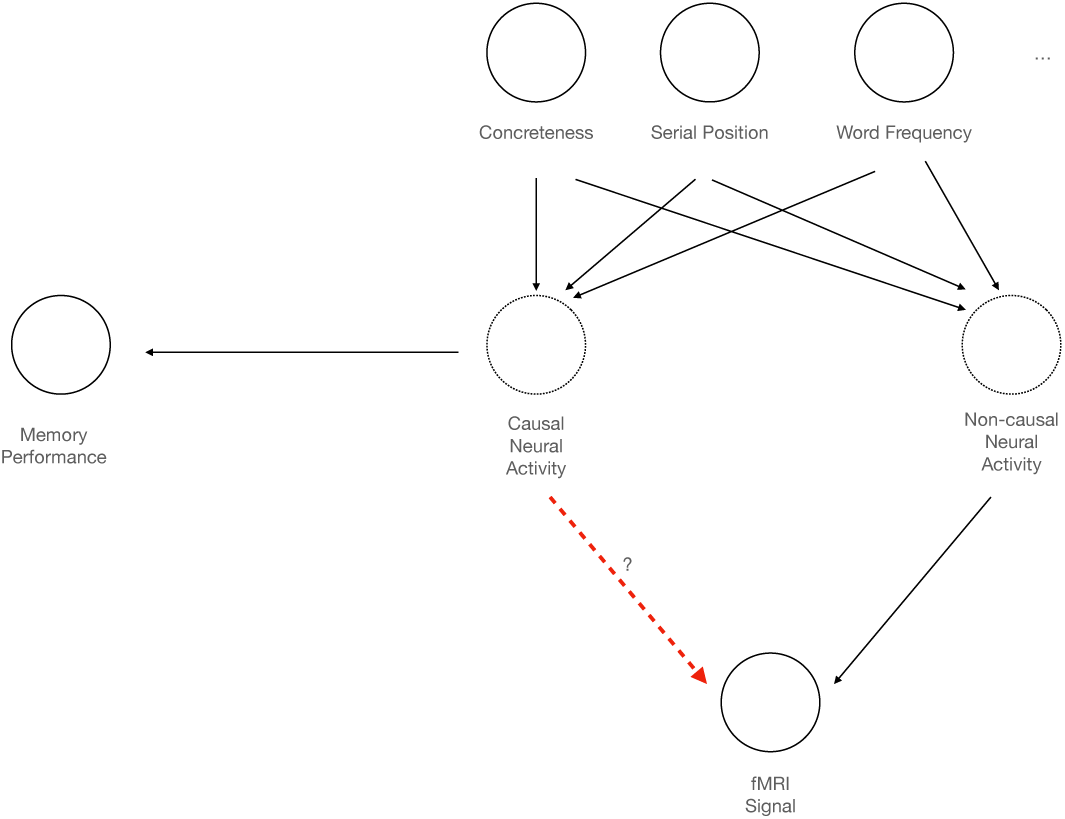
A directed acyclic graph representing a causal for memory if exogenous variables were the only variables that influenced memory. Solid circles indicate observed variables while dotted circles indicate unobserved.

**Figure S3:**
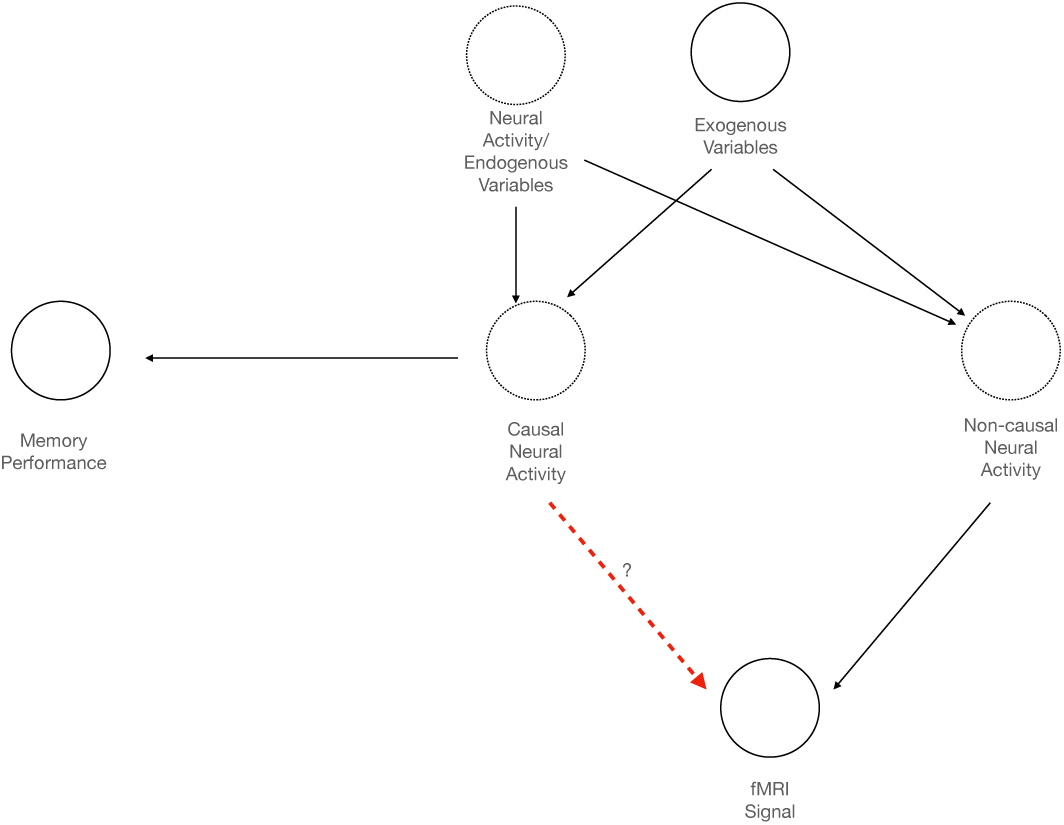
A graph including all endogenous and exogenous variables that affect memory.

**Figure S4:**
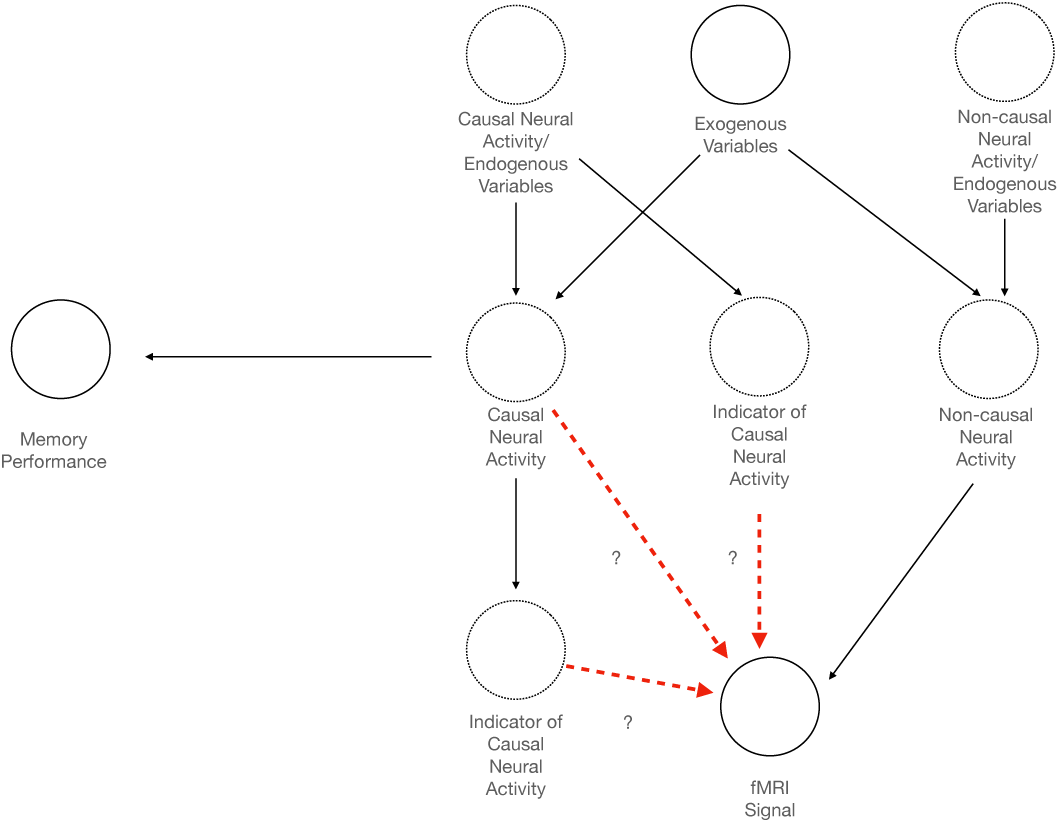
A graph showing the distinction between causal encoding activity and indicators of causal encoding activity. Using our approach, we cannot distinguish between these two types of variables, due to the fact that the endogenous variables are uncontrolled.

## Footnotes

1 We do not hold neural activity constant because for most signals, some neural activity is downstream and therefore would be a post-treatment variable which might confound causal identification. (Montgomery et al., 2018; Rosenbaum, 1984)

2 Descriptions of components of models used in Figure 5 and 6 are indicated in the main text using bold font

3 This was the smallest possible p-value in our permutation test given that we ran 500 samples. The true *p*-value may be much smaller but we do not have the precision to report it.

4 https://afni.nimh.nih.gov/pub/dist/doc/program_help/3dBlurToFWHM.html

5 https://github.com/ThomasYeoLab/CBIG/tree/master/stable_projects/brain_parcellation/Schaefer2018_LocalGlobal)

6 https://neurovault.org/collections/3731/; https://identifiers.org/neurovault.collection:3731

7 ^7^https://www.fil.ion.ucl.ac.uk/spm/software/spm12/

## Notes

### Competing Interest Statement

The authors have declared no competing interest.

